# Grant Review Feedback: Appropriateness and Usefulness

**DOI:** 10.1101/2020.11.24.396192

**Authors:** Stephen Gallo, Karen Schmaling, Lisa Thompson, Scott Glisson

## Abstract

The primary goal of the peer review of research grant proposals is to evaluate their quality for the funding agency. An important secondary goal is to provide constructive feedback to applicants for their resubmissions. However, little is known about whether review feedback achieves this goal. In this paper, we present a mixed methods analysis of responses from grant applicants regarding their perceptions of the effectiveness and appropriateness of peer review feedback they received from grant submissions. Overall, 56%-60% of applicants determined the feedback to be appropriate (fair, well-written, and well-informed), although their judgments were more favorable if their recent application was funded. Importantly, independent of funding success, women found the feedback better written than men, and more white applicants found the feedback to be fair than non-white applicants. Also, perceptions of a variety of biases were specifically reported in respondents’ feedback. Less than 40% of applicants found the feedback to be very useful in informing their research and improving grantsmanship and future submissions. Further, negative perceptions of the appropriateness of review feedback were positively correlated with more negative perceptions of feedback usefulness. Importantly, respondents suggested that highly competitive funding pay-lines and poor inter-panel reliability limited the usefulness of review feedback. Overall, these results suggest that more effort is needed to ensure that appropriate and useful feedback is provided to all applicants, bolstering the equity of the review process and likely improving the quality of resubmitted proposals.

## Introduction

Most scientific research funding agencies utilize a peer review system to evaluate submitted projects and select the most meritorious for funding. At the National Institutes of Health (NIH), after a grant application is peer reviewed, the scores and comments from the reviewers are sent back to the applicant (NIH 2018). If it is not funded, the applicant must address the reviewer comments if they are to resubmit an updated version of their application (NIH 2020 and NIAID 2020). NIH reviewers evaluate the scientific and technical merit of each proposal and are required to justify their scores with comments on the proposal’s strengths and weaknesses. The comments “should be clear enough that an investigator has a sense of what needs to be done in order to craft a more competitive application if the current version is unfunded” (NIH CSR 2020). Thus, although not listed as a core value of the NIH peer review system, reviewer feedback to applicants for the purposes of improving investigator grantsmanship and the overall quality of applications is an important, if secondary, purpose of grant peer review (NIH 2019). Little empirical data exist that document whether grant review feedback is effective in informing applicants and improving applications.

Useful feedback is likely particularly important to help improve resubmitted applications, which are proportionally more likely to be funded than new applications, although funding rates are low overall (Lauer 2016, Lauer 2017, Haggerty and Fenton 2018). The success of resubmitted applications depends on an applicant’s ability to address reviewer concerns in their resubmissions (NIAID 2020). Resources are available to guide scientists as to the best approaches (Boss and Eckert 2003, Sutcivni 2017). However, while reviewers tend to believe their feedback is useful to move scientific fields forward (Irwin et al. 2013), it is unclear if applicants find the reviewer comments useful for resubmission. In 2015, NIH surveyed all stakeholders of their peer review process to learn about perceptions of the peer review process, including the value of reviewer feedback by applicants (NIH 2017). The results reveal only 53% of applicants “strongly agreed/agreed that the information in their summary statement helped them to focus on problem areas in the application” (NIH 2017, p.4). However, this result still does not indicate whether the feedback was specifically useful; we could define usefulness of feedback in terms of specifically improving resubmissions, and generally improving applicant grantsmanship and informing future scientific endeavors. Importantly, the responses to the NIH survey were greatly influenced by whether the applicants were recently funded, where “favorable responses were more prevalent among funded applicants than applicants whose applications were not funded” (NIH 2017, p.17).

The score an initial grant receives (including whether or not it was triaged) is a key predictor of whether the applicant decides to resubmit (Boyington et al. 2016, Lauer 2017), but it is not clear if applicant perception of the review feedback is also a significant predictor of resubmission. Recent studies have suggested that women applicants interpret peer review feedback more negatively than men, which was associated with reduced motivation to resubmit (Biernat et al. 2020). It is well documented that women submit fewer research applications than men (Hechtman et al. 2018) and that under-represented minority (URM) women of color submit even fewer, have lower funding success compared to white women, and are less likely to resubmit unfunded applications (Ginther et al. 2016). Specifically, African American scientists have been found to submit fewer new applications, fewer resubmissions, and are funded at a lower rate than white scientists (Ginther et al. 2011, Mervis 2016). Thus, a negative interpretation of feedback by URM scientists compared to majority group scientists could reduce the rate of resubmissions, which may contribute to funding disparities. Despite this possibility, little is known about how applicants view the usefulness of feedback in guiding their resubmissions.

In addition, it is unknown whether applicants deem the feedback they receive to be appropriate, which we could define as being well-written, unbiased and based on expert perspective. Although the peer review process may be subject to a variety of biases (Lee et al., 2013), it is unclear if applicants perceive these biases to be explicitly present in the review feedback. Some research has suggested that review feedback to women applicants contains different language than to men, particularly in the evaluation of the investigator’s leadership abilities (Pier et al. 2018). In the 2015 NIH survey, only 54% of applicants perceived the peer review process as fair. In addition, 91% of funded applicants and 53% of unfunded applicants “agreed that their application was evaluated by reviewers with the appropriate expertise,” suggesting that funding success influenced perceptions of the appropriateness of review feedback. However, little is known about the influence of applicant demographics on perceptions of the appropriateness of review feedback and if there is any association between perceived appropriateness and perceived usefulness of the feedback for resubmission. The objective evaluation of reviewer comments is made difficult by a lack of access to critiques of unfunded applications (Gurwitz et al. 2014, Gropp et al. 2017).

Given the paucity of data surrounding grant review feedback, particularly from an applicant’s perspective, we created a survey for research scientists that asked applicant respondents to rate and comment on the review feedback they received on their most recent submission. Three publications have resulted from this survey (Gallo et al. 2018, Gallo et al. 2020a, Gallo et al. 2020b), but none of them focused on the questions related to the usefulness and appropriateness of peer review feedback; these questions are now addressed in the proceeding analysis.

## Methods

### Survey Construction and Mixed Methods Approach

To our knowledge, there are no reports in the peer-reviewed literature that have queried grant applicants about their perceptions of the peer review process. Therefore, the authors brainstormed questions to address the range of relevant content associated with peer review feedback. These ideas were also informed by input from scientists, scientific review administrators and research funders, and the literature on peer review processes. A draft survey was examined by the authors and others experienced in peer review for its clarity, face validity, and coverage of relevant content.

The survey included questions on applicant perceptions of review feedback that yielded nominal and ordinal quantitative data and open text fields at the end of every section that yielded qualitative data. Our rationale for using this mixed methods approach (Figure 1) was to improve our analysis of applicant perceptions of the appropriateness and usefulness of review feedback. Text was associated with specific questions with quantitative answer options, as described below, and then used to elaborate on respondents’ quantitative answers.

**Figure 1.**
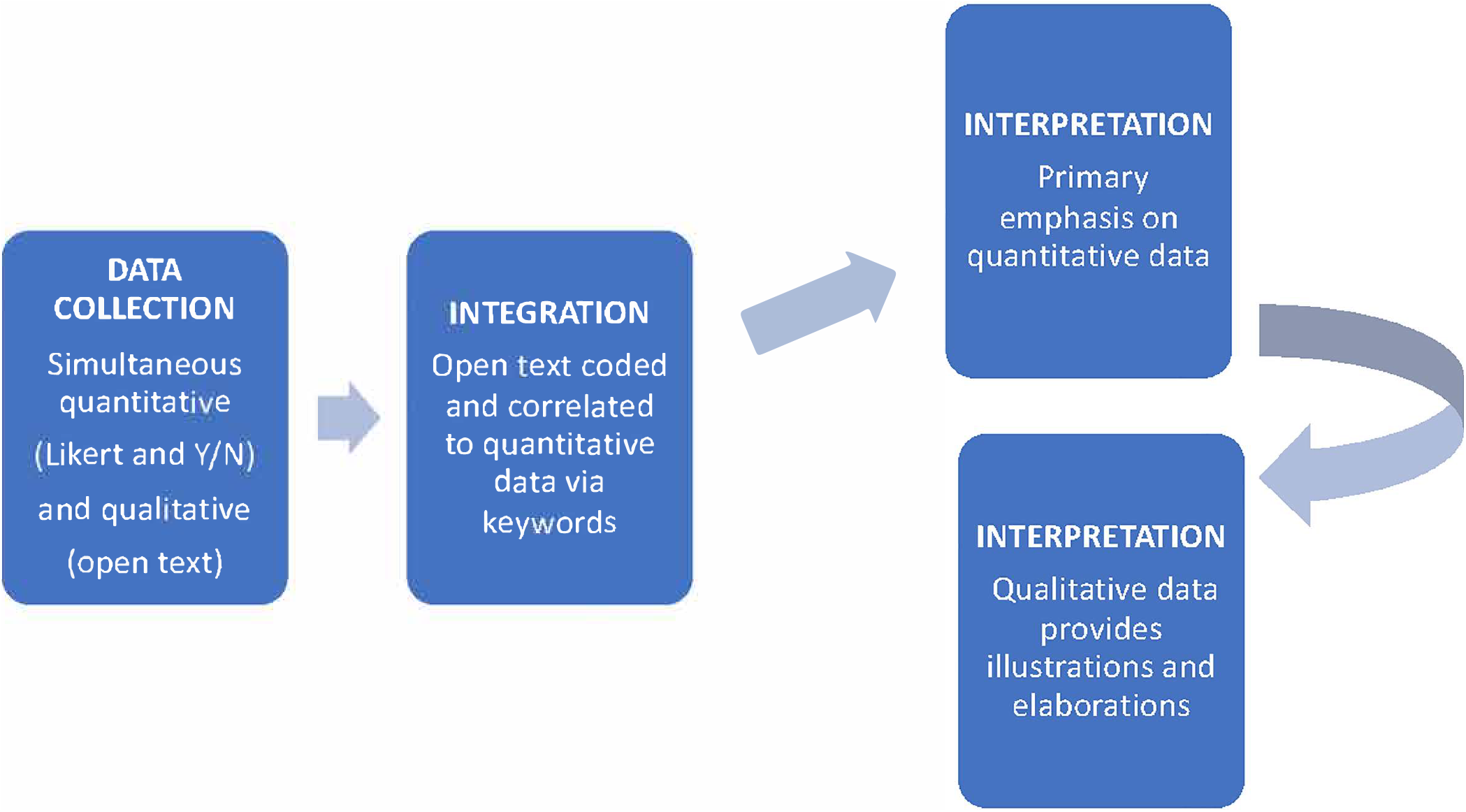
Schematic of our mixed methods approach.

### Survey Data Collection

The study was reviewed by the Washington State University Office of Research Assurances (Assurance# FWA00002946) and granted an exemption from IRB review consistent with 45 CFR 46.101(b)(2). Respondents were fully informed about the purpose, importance, intended use of the survey, and how we acquired their email address, and consented by their participation. The general survey description has been included in previous publications (Gallo et al. 2018); the survey contained 60 questions in 5 subsections, three of which were included in this analysis (Section 1: Demographics, Section 2: Grant submission and peer review experience, and Section 3: Investigator attitudes toward grant review). Specifically, in this analysis we examined questions related to the usefulness and appropriateness of review feedback, from an applicant perspective. As suggested above, we define usefulness of feedback in terms of: improving future resubmissions; improving applicant grantsmanship; and informing future scientific endeavors. We define appropriateness of feedback as being: well-written, cohesive and balanced; fair and unbiased; and based on a relevant and expert perspective.

Questions related to usefulness had Likert ordinal responses, questions related to appropriateness had nominal (yes/no) responses, and there was an open text field for respondent comments at the end of this survey section. Respondents could choose to select no answer/prefer not to answer, and comments were not compulsory. The full survey is listed in other publications and is included as a supplement to this manuscript (Gallo et al. 2018, File S1).

The survey was emailed in September 2016 to a total of 13,091 scientists in the American Institute for Biological Sciences’ database, which was developed for assisting in the recruitment of scientific expert reviewers to evaluate biomedical research applications for several funding organizations. Scientists recruited for this survey work in the biomedical area, ranging from clinical psychology to molecular biology. Of the individuals invited to participate in the survey, 74% had applied for research funding in the last 3 years. Reminder emails were sent out to non-responders two weeks before the survey was closed.

### Data Summarization

The survey was open for two months, with a reminder sent 2 weeks before the survey closed. Responses were exported and Stat Plus software was used for the analysis. In this study, respondents were included if they completed the entire survey and answered three questions affirmatively (or greater than one): 2a (Have you submitted a grant for peer review in the last 3 years?), 2b (How many grant applications have you submitted in that time frame?), and 2c (Did you receive reviewer feedback on your last grant submission?).

For the quantitative data, medians and percentages were calculated for the responses to the survey questions of interest, standard 95% confidence intervals were reported for the ordinal responses, and binomial proportion confidence intervals for the proportion data. To test the internal consistency of the survey’s quantitative items, we calculated Cronbach’s alpha for the four questions related to Usefulness (Q4-7), which yielded an alpha of 0.87. The initial assessment of demographic factors related to responses was conducted through multiple binary logistic (for appropriateness responses) and ordinal logistic (for usefulness responses) regression for nominal and ordinal responses, respectively. Nominal responses were coded as Yes=1, No=0 and Likert responses were on a scale of 1 to 5 with 1 as the most useful and 5 as the least useful. Demographic data variables of applicant characteristics were categorical in nature, and were coded as 0 or 1; where there were unequal distributions between categories, the more frequently appearing category was coded as a 1. These variables included race (coded as 1=white, 0=non-white), gender (man=1, 0=woman), career stage (early/mid-career=1, late stage career/emeritus=0), age (50 and over=1, under 50=0), degree (PhD=1, non-PhD=0), and funding status (recently funded=1, unfunded=0). For the regression analyses, predictors were entered together as a block. Chi square and Mann Whitney tests were used for post-hoc comparisons, using phi coefficient or the Z-score/√N, respectively, as the effect size. Differences between groups were determined to be significant if there was no overlap in confidence interval and tests for differences yielded p-values <0.01. Variance inflation factors were used to examine for potential multicollinearity of the review feedback and demographic variables.

For the qualitative data, respondents’ comments from Section 3 of the survey (investigator attitudes toward grant review) were extracted (N=216). Comments were not mandatory, so the demographic characteristics of those who did and did not comment were compared. Comments were coded for content related to two groups of survey questions on the appropriateness (Q1-3) and usefulness (Q4-7) of the review feedback (Supplementary Table 1). These quotes were sorted into sub-categories by the presence of keywords that were related to the survey questions. Specifically, these keywords were “written” for Q1, “bias” for Q2, and “expertise” for Q3. In general, we chose these keywords for Q1-3 because they most directly addressed the survey questions and allowed for the most unambiguous and objective coding to specific questions. However, because several keywords in questions 4-7 appeared infrequently in the comments (namely “grantsmanship,” scientific endeavor” or “research area”), we used the keyword “useful” and its synonym “helpful” to identify quotes related to Q4-7. Keyword searches utilized the simple word matching function in Excel and included all instances of the base words including suffix usage (e.g. usefulness) and different tenses (e.g. biased). Quotes could be sorted into multiple groups if more than one keyword was present. A total of 79 quotes (37% of all quotes) was explicitly grouped in this way, with 19% of those quotes being represented in more than one group (Supplementary Table 1). Although many of the ungrouped quotes had some relevance to the questions asked in the survey, only explicitly grouped quotes were included in the analysis below.

Although the survey questions were worded positively, the associated comments could be of positive or negative valence. For example, both “reviewers had the expertise” and “reviews were unfair, biased, and lacked appropriate expertise” would be attributed to question Q3 (“Based on the reviewer feedback you received, do you feel that the reviewers had the appropriate expertise to evaluate your grant application?”) and reflected both positive and negative valence, respectively. Thus, each comment was also coded for valence: positive, negative, or both positive and negative (all of the coded comments had some positive or negative valence) (Supplementary Table 1). Two authors coded all comments for valence; there was agreement for 90% of comments and the 10% with disagreements were discussed until consensus was achieved.

## Results

### Response Rate and Demographics

Of the 13,091 individuals contacted for this survey, 1231 responded, for a 9.4% response rate. Of the 1231 respondents, 634 answered questions 2a, 2b, and 2c in the affirmative, indicating they had submitted a proposal in the last 3 years and received review feedback. The remaining results focus on this sample of 634 participants. Over half of the responses were collected within a few hours of sending the emailed invitations to complete the survey. Another, smaller wave of responses was collected after a reminder email was sent. Comparisons of the quantitative answers to questions for Usefulness and Appropriateness were nearly identical for these two groups (Supplementary Table 2).

Sample demographics are listed in Table 1: the majority of respondents were men, Caucasian, academic PhDs in a late career stage. They had submitted a median of 5.0 ± 0.1 applications in the last 3 years. The overall funding success rate was 40% (95%CI, 36% to 44%), which did not vary significantly by gender, race, age, career stage, degree, and organization (Table 1).

**Table 1.** Demographics and Success Rates

| Factor (N=634) |  | Proportion (N) | % Funding Success |
| --- | --- | --- | --- |
| Gender | Men | 63% (398) | 41% |
|  | Women | 37% (236) | 39% |
| Age | Under 50 | 29% (177) | 37% |
|  | 50 and Over | 71% (436) | 43% |
| Race/Ethnicity | White | 75% (478) | 40% |
|  | Non-White | 25% (156) | 41% |
| Degree Type | PhD | 83% (525) | 40% |
|  | Non-PhD | 17% (109) | 43% |
| Organization | Academia | 88% (552) | 39% |
|  | Non-Academia | 12% (72) | 50% |
| Career Stage | Early | 4% (23) | 30% |
|  | Mid | 31% (191) | 39% |
|  | Late/Tenured | 62% (385) | 44% |
|  | Emeritus | 4% (27) | 26% |

Some differences were evident in the demographics of respondents who made a comment versus non-commenting respondents (Supplementary Table 3). Non-whites represented only 17% (95%CI [12% to 22%]) of the commenting respondent pool compared to representing 29% (95%CI [26% to 34%]) of the non-commenting respondent pool (X^2^ [1]=11.2, p=0.0008, phi=0.13). Additionally, respondents who made a comment were more likely to be older and in later career stages than non-respondents (Supplementary Table 3). Other demographic variables -- degree, organization and gender -- did not differ between commenting and non-commenting groups. However, it was also noted that 32% (N=67; CI [26% to 38%]) of respondents who made a comment reported being recently funded compared to 44% (N=183; 95%CI [39% to 49%]) of respondents who did not make a comment (X^2^ [1]=8.9, p=0.0029, phi=0.12).

Although the survey did not ask respondents to which funding agency they had applied, we did search the comments for mention of the two largest US funders, NIH and NSF. Of the comments that mentioned funding agency, 14% mentioned NSF and 86% mentioned NIH.

### Overview of Mixed Method Results of Respondent Comments and Ratings

The sections below present the results of the qualitative analysis of the comments and the quantitative ratings of the associated survey questions Q1-7 (Supplementary Table 1). The 13 quotes listed in Supplementary Table 4 were specifically chosen as examples that captured a particular theme associated with the questions asked in the survey. It should be noted, though, that respondents’ comments often had a negative valence and respondents with comments tended to be more negative even in the quantitative portions of the survey as compared to respondents without comments (Supplementary Table 5). We then examine how these results vary with applicant demographics, and the relationship between responses related to appropriateness and usefulness.

### Appropriateness of Feedback

Overall, only 56% (95%CI [52% to 60%]) of respondents thought grant review feedback was well written, cohesive, and balanced. Respondents indicated issues related to the structure and length of the feedback (Supplementary Table 4; Q1.1) and that often reviewers do not support their score with comments (Supplementary Table 4; Q1.2).

Additionally, 60% (95%CI [56% to 64%]) of respondents perceived grant reviewer feedback as fair and unbiased. Comments, however, identify various types of perceived biases toward specific application content, including bias against topic areas, bias against innovation, and methodology/model bias (Supplementary Table 4; Q2.1). Some comments also specifically mentioned biases against the applicants (Supplementary Table 4; Q2.2 and Q2.3). Some respondents suggested the impact biased reviews can have, particularly in an era of low funding success rates (Supplementary Table 4; Q2.4).

In terms of reviewer qualifications, 58% (95%CI [54% to 62%]) of respondents judged the reviewers to have appropriate expertise to evaluate their grant application, based on the reviewer feedback they received. In their comments, respondents identified a lack of reviewer expertise for interdisciplinary proposals, clinical proposals, statistical portions of the proposals, proposals with different animal models, and a lack of expertise in specific areas of science (Supplementary Table 4; Q3.1 and Q3.2).

### Usefulness of Feedback

Overall, only 38% (median of 3.0; 95%CI [2.9 to 3.1]) found the grant review feedback they received on their last grant submission to be mostly useful or very useful. Further, only 30% (median of 3.0; 95%CI [2.9 to 3.1]) thought it was mostly useful or very useful in improving their grantsmanship; only 35% (median of 3.0; 95%CI [2.9 to 3.1]) found the review feedback was mostly useful or very useful in improving their future submissions; and only 26% (median value of 3.0; 95%CI [2.9 to 3.1]) felt the feedback was mostly or very useful in informing their future scientific endeavors in the proposed research area. Based on these data, the majority of applicants did not find the reviewer feedback to be highly useful.

Few comments specifically mentioned the usefulness of the feedback in terms of grantsmanship (Q5) or future scientific endeavors (Q7); more were related to the usefulness of the feedback in improving future submissions (Q6). Some remarked on how they received constructive criticism that helped improve their applications (Supplementary Table 4; Q4-7.1). However, some remarked that inconsistent feedback from different sets of reviewers evaluating resubmissions reduces usefulness (Supplementary Table 4; Q4-7.2). Others commented that usefulness is hampered by a perceived lack of expertise (Supplementary Table 4; Q4-7.3). Several comments mentioned that the feedback format and lack of suggestions for improvement limit usefulness (Supplementary Table 4; Q4-7.4). Finally, some mentioned that usefulness of feedback was ultimately restricted by funding success rates (Supplementary Table 4; Q4-7.5).

### Perceptions of Feedback and Demographics

We used multiple regression analysis to examine the effects of demographic variables on perceived appropriateness and usefulness of feedback. As seen in Supplementary Table 6, the variance inflation factors between most of these demographic variables is low.

We first analyzed the relationships between demographic variables and the nominal responses related to the appropriateness of review feedback using binary logistic regression. We found significant differences in terms of funding status for responses to all three questions related to appropriateness, as indicated by the reported odds ratios (Table 2).

**Table 2.**
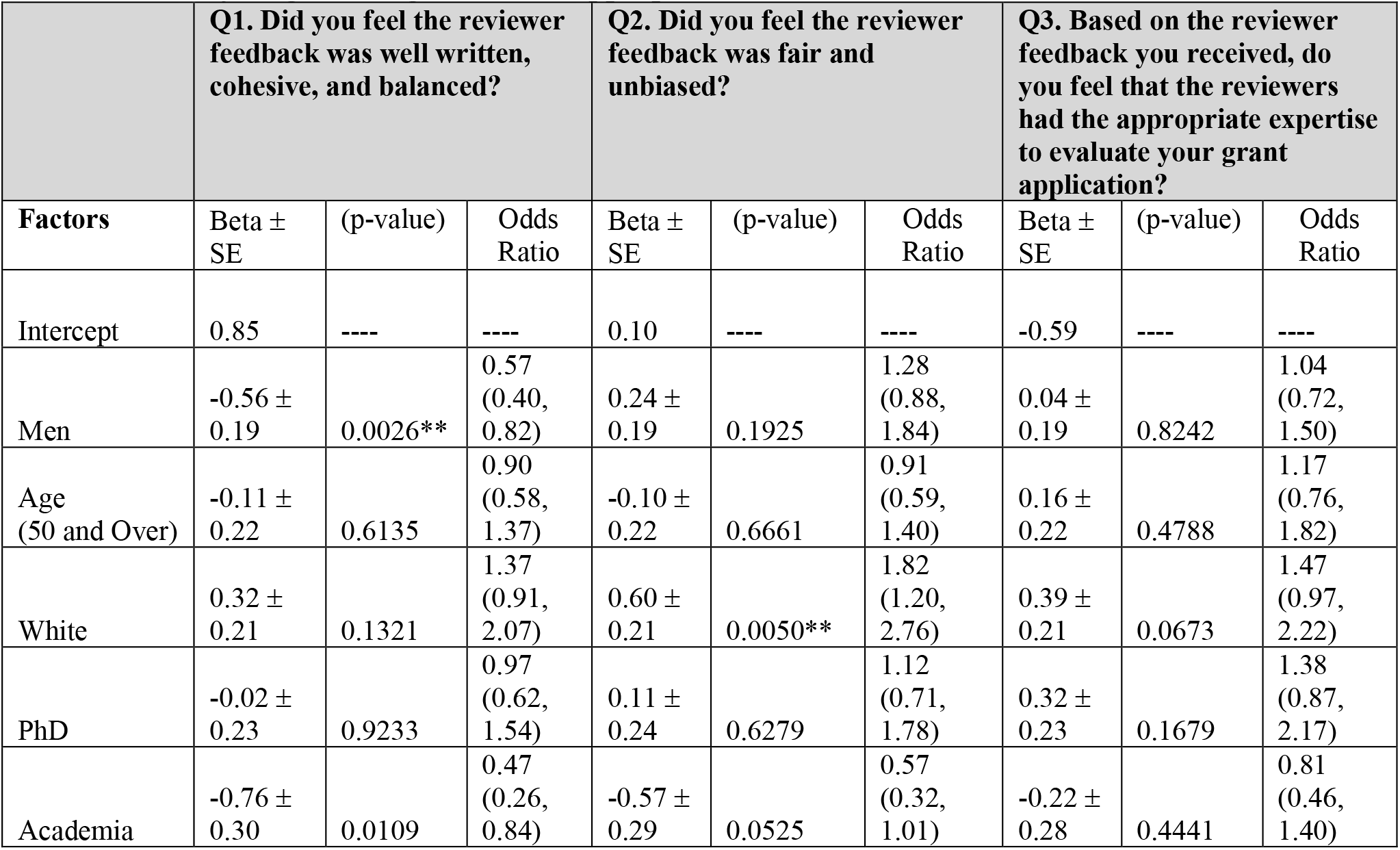
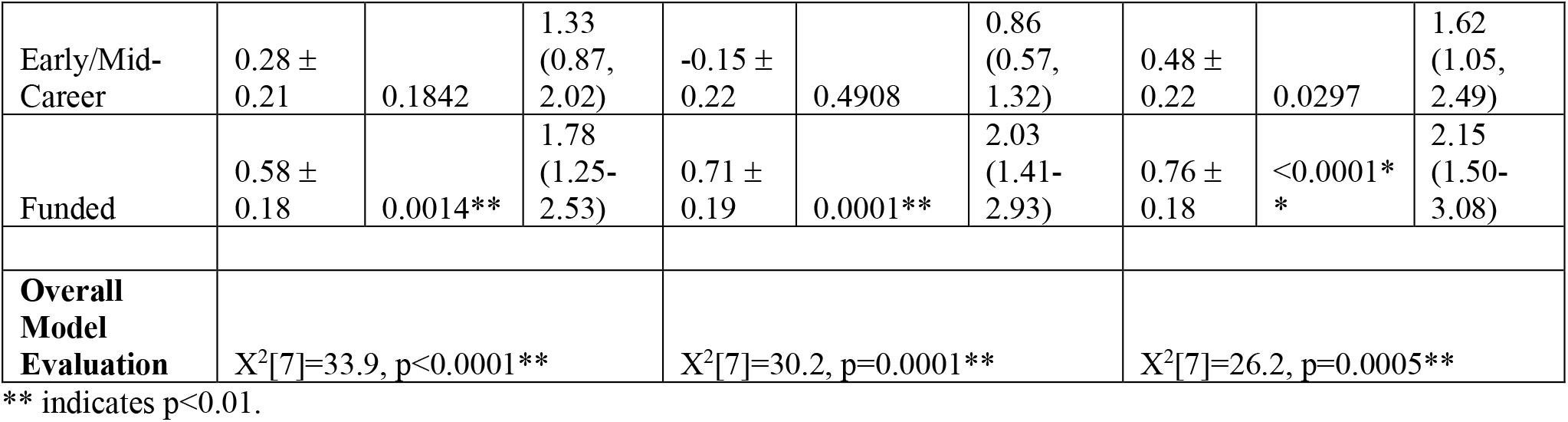
Binary Logistic Regression of Appropriateness of Review Feedback

|  | **Q1. Did you feel the reviewer feedback was well written, cohesive, and balanced?** |  |  | **Q2. Did you feel the reviewer feedback was fair and unbiased?** |  |  | **Q3. Based on the reviewer feedback you received, do you feel that the reviewers had the appropriate expertise to evaluate your grant application?** |  |  |
| --- | --- | --- | --- | --- | --- | --- | --- | --- | --- |
| **Factors** | Beta ± SE | (p-value) | Odds Ratio | Beta ± SE | (p-value) | Odds Ratio | Beta ± SE | (p-value) | Odds Ratio |
| Intercept | 0.85 | ---- | ---- | 0.10 | ---- | ---- | -0.59 | ---- | ---- |
| Men | -0.56 ± 0.19 | 0.0026** | 0.57 (0.40, 0.82) | 0.24 ± 0.19 | 0.1925 | 1.28 (0.88, 1.84) | 0.04 ± 0.19 | 0.8242 | 1.04 (0.72, 1.50) |
| Age (50 and Over) | -0.11 ± 0.22 | 0.6135 | 0.90 (0.58, 1.37) | -0.10 ± 0.22 | 0.6661 | 0.91 (0.59, 1.40) | 0.16 ± 0.22 | 0.4788 | 1.17 (0.76, 1.82) |
| White | 0.32 ± 0.21 | 0.1321 | 1.37 (0.91, 2.07) | 0.60 ± 0.21 | 0.0050** | 1.82 (1.20, 2.76) | 0.39 ± 0.21 | 0.0673 | 1.47 (0.97, 2.22) |
| PhD | -0.02 ± 0.23 | 0.9233 | 0.97 (0.62, 1.54) | 0.11 ± 0.24 | 0.6279 | 1.12 (0.71, 1.78) | 0.32 ± 0.23 | 0.1679 | 1.38 (0.87, 2.17) |
| Academia | -0.76 ± 0.30 | 0.0109 | 0.47 (0.26, 0.84) | -0.57 ± 0.29 | 0.0525 | 0.57 (0.32, 1.01) | -0.22 ± 0.28 | 0.4441 | 0.81 (0.46, 1.40) |
| Early/Mid-Career | 0.28 ± 0.21 | 0.1842 | 1.33 (0.87, 2.02) | -0.15 ± 0.22 | 0.4908 | 0.86 (0.57, 1.32) | 0.48 ± 0.22 | 0.0297 | 1.62 (1.05, 2.49) |
| Funded | 0.58 ± 0.18 | 0.0014** | 1.78 (1.25-2.53) | 0.71 ± 0.19 | 0.0001** | 2.03 (1.41-2.93) | 0.76 ± 0.18 | <0.0001* | 2.15 (1.50-3.08) |
| **Overall Model Evaluation** | X<sup>2</sup>[7]=33.9, p<0.0001** |  |  | X<sup>2</sup>[7]=30.2, p=0.0001** |  |  | X<sup>2</sup>[7]=26.2, p=0.0005** |  |  |

\*\* indicates p<0.01.

For example, for the Q1 regression, the factor of funding status had an odds ratio of 1.78 (95% CI, 1.25 to 2.53). Thus, respondents who were funded were nearly twice as likely to indicate that the review feedback was well-written, cohesive, and balanced as compared to respondents who were not funded; indeed, 63% (95%CI, 57% to 69%) of funded respondents found the feedback to be well-written, cohesive and balanced compared to 51% (95%CI, 46% to 56%) of unfunded respondents (X2 [1]=9.2, p=0.0024, phi=0.12). Similarly, funded respondents were more likely to find the feedback was fair and unbiased (Q2 Table 2): 71% (95%CI, 65% to 77%) of funded respondents versus 53% (95%CI, 48% to 58%) of unfunded respondents (X^2^ [1]=18.0, p<0.0001, phi=0.18). A greater number of funded respondents perceived the reviewers to have appropriate expertise to evaluate their proposal (Q3; Table 2): 68% (95%CI, 62% to 74%) of funded respondents versus 51% (95%CI, 46% to 56%) of unfunded respondents (X^2^ [1]=17.0, p<0.0001, phi=0.17).

No differences were observed by career stage, age, organization or degree with respect to perceptions of appropriateness (Table 2). However, gender and race were found to predictperceptions of the appropriateness of review feedback (Table 2). Women were significantly more likely to rate the reviewer feedback as well written, cohesive, and balanced compared than men (64% [95%CI, 58% to 70%] and 53% [95%CI, 48% to 58%], respectively) (X^2^ [1]=9.3, p=0.0023, phi=0.12). Whites were significantly more likely to rate the feedback as fair and unbiased than non-whites ((64%, 95%CI [60% to 68%]) and 49%, 95%CI [41% to 57%], respectively) (X^2^ [1]=9.2, p=0.0024, phi=0.12). These differences were not due to funding success, as the rates were similar between groups (Table 1). In terms of reviewer expertise, responses to Q3 did not vary significantly by race or gender (Q3 Table 2).

Overall, for the responses related to the appropriateness of review feedback, no thematic differences were found between the comments made by non-white versus white applicants. Similarly, no thematic differences were found between the comments made by women versus men.

We then analyzed the relationships between demographic variables and the ordinal Likert responses (1-5 where 1 is most useful) related to the usefulness of review feedback using multiple ordinal logistic regression. None of the regression models for responses related to questions of general usefulness (Q4), usefulness in improving grantsmanship (Q5), usefulness in improving future submissions (Q6), and usefulness in informing future scientific endeavors (Q7) were found to explain significant proportions of the variance in the responses (Table 3). Moreover, none of the funding and demographic factors (including race, gender, career stage, age, degree, or organization) were found to be significant predictors of these responses.

**Table 3.** Ordinal Logistic Regression of Usefulness of Review Feedback

|  | Q4. On a scale of 1-5 (1 most useful, 5 least useful), overall how useful was the reviewer feedback you received on your last grant submission? |  |  | Q5. On a scale of 1-5 (1 most useful, 5 least useful), how useful was the reviewer feedback in improving your grantsmanship? |  |  | Q6. If you were not funded, on a scale of 1-5 (1 most useful, 5 least useful), how useful was the reviewer feedback in improving your future submissions? |  |  | Q7. On a scale of 1-5 (1 most useful, 5 least useful), how useful was the reviewer feedback in informing your future scientific endeavors in the proposed research area? |  |  |
| --- | --- | --- | --- | --- | --- | --- | --- | --- | --- | --- | --- | --- |
| Factors | Beta ± SE | p-value | Odds Ratio | Beta ± SE | p-value | Odds Ratio | Beta ± SE | p-value | Odds Ratio | Beta ± SE | p-value | Odds Ratio |
| Intercept 1/2 | -2.78 ± 0.40 | <0.0001** | n.a. | -2.40 ± 0.40 | <0.0001** | n.a. | -2.95 ± 0.43 | <0.0001** | n.a. | -2.82 ± 0.41 | <0.0001** | n.a. |
| Intercept 2/3 | -0.64 ± 0.38 | 0.0903 | n.a. | -0.57 ± 0.37 | 0.1250 | n.a. | -1.21 ± 0.40 | 0.0026** | n.a. | -1.13 ± 0.39 | 0.0035** | n.a. |
| Intercept 3/4 | 0.56 ± 0.38 | 0.1390 | n.a. | 0.56 ± 0.37 | 0.1292 | n.a. | -0.15 ± 0.40 | 0.7042 | n.a. | 0.14 ± 0.38 | 0.7141 | n.a. |
| Intercept 4/5 | 1.97 ± 0.39 | <0.0001** | n.a. | 1.58 ± 0.38 | <0.0001** | n.a. | 1.15 ± 0.40 | 0.0042** | n.a. | 1.16 ± 0.38 | 0.0025** | n.a. |
| Men | -0.04 ± 0.16 | 0.8051 | 0.96 (0.71, 1.30) | 0.07 ± 0.16 | 0.6718 | 1.07 (0.79, 1.45) | -0.26 ± 0.17 | 0.1262 | 0.77 (0.55, 1.08) | 0.19 ± 0.16 | 0.2313 | 1.21 (0.89, 1.65) |
| Age (50 and over) | -0.02 ± 0.19 | 0.9037 | 0.98 (0.68, 1.41) | 0.17 ± 0.18 | 0.3605 | 1.18 (0.83, 1.68) | 0.12 ± 0.19 | 0.5411 | 1.13 (0.77, 1.64) | 0.08 ± 0.19 | 0.6477 | 1.09 (0.76, 1.57) |
| White | -0.07 ± 0.18 | 0.7141 | 0.94 (0.66, 1.33) | 0.26 ± 0.18 | 0.1411 | 1.30 (0.92, 1.85) | -0.07 ± 0.19 | 0.6982 | 0.93 (0.64, 1.34) | 0.06 ± 0.18 | 0.7330 | 1.06 (0.75, 1.51) |
| PhD | -0.13 ± 0.20 | 0.5124 | 0.88 (0.59, 1.30) | 0.19 ± 0.20 | 0.3326 | 1.21 (0.82, 1.78) | -0.26 ± 0.22 | 0.2364 | 0.77 (0.51, 1.18) | -0.19 ± 0.20 | 0.3541 | 0.83 (0.56, 1.23) |
| Academia | 0.28 ± 0.24 | 0.2495 | 1.32 (0.82, 2.11) | -0.07 ± 0.23 | 0.7501 | 0.93 (0.58, 1.47) | -0.10 ± 0.26 | 0.4031 | 0.90 (0.54, 1.49) | -0.03 ± 0.24 | 0.8947 | 0.97 (0.61, 1.55) |
| Early/Mid-Career | -0.20 ± 0.18 | 0.2630 | 0.82 (0.57, 1.16) | -0.04 ± 0.18 | 0.8417 | 0.97 (0.68, 1.36) | -0.25 ± 0.19 | 0.1936 | 0.78 (0.54, 1.13) | -0.04 ± 0.18 | 0.8137 | 0.96 (0.67, 1.37) |
| Funded | -0.30 ± 0.15 | 0.0473 | 0.73 (0.54, 1.00) | -0.28 ± 0.15 | 0.0619 | 0.75 (0.56, 1.01) | -0.14 ± 0.18 | 0.0042** | 0.87 (0.61, 1.25) | -0.21 ± 0.16 | 0.1725 | 0.81 (0.60, 1.10) |
| **Overall Model Evaluation** | X<sup>2</sup>[7]=6.9, p=0.4361 |  |  | X<sup>2</sup>[7]=8.3, p=0.3101 |  |  | X<sup>2</sup>[7]=7.0, p=0.4268 |  |  | X<sup>2</sup>[7]=5.0, p=0.6596 |  |  |

\*\* indicates p<0.01

### Appropriateness versus Usefulness of Feedback

Perceived usefulness of review feedback may be associated with grant resubmission rates, but the associations of perceived appropriateness of feedback and applicant behavior are unclear. Based on our results that the majority of respondents did not find review feedback useful and a large minority of respondents did not find the feedback appropriate, it is likely that applicants who don’t find the feedback appropriate also don’t find it useful. In fact, some comments in our survey suggested usefulness was limited by the lack of appropriate expertise. To test this assumption, we compared respondents who found the review feedback they received to be fair and unbiased (Q2) to respondents who did not. For these two groups, we examined their answers to the questions concerning the usefulness of the feedback (Q4-Q7). This comparison of usefulness and appropriateness is listed in Table 4. The results revealed significantly more negative perceptions of the usefulness of the feedback for those who also felt the feedback was biased compared to those who felt the feedback was unbiased.

**Table 4.** Review Feedback Appropriateness versus Usefulness

| Question | Q2. Feedback was fair and unbiased (Yes) |  | Q2. Feedback was unfair and biased (No) |  | Fair versus Unfair |
| --- | --- | --- | --- | --- | --- |
| | Median ± 95%CI | N | Median ± 95%CI | N | Mann-Whitney |
| Q4. On a scale of 1-5 (1 most useful, 5 least useful), overall how useful was the reviewer feedback you received on your last grant submission? | 2.0 ± 0.10 | 356 | 4.0 ± 0.14 | 234 | U[234, 356]=61,212, p<0.0001**, r=0.40 |
| Q5. On a scale of 1-5 (1 most useful, 5 least useful), how useful was the reviewer feedback in improving your grantsmanship? | 3.0 ± 0.12 | 358 | 4.0 ± 0.15 | 233 | U[233, 358]=57,945, p<0.0001**, r=0.33 |
| Q6. If you were not funded, on a scale of 1-5 (1 most useful, 5 least useful), how useful was the reviewer feedback in improving your future submissions? | 3.0 ± 0.13 | 271 | 4.0 ± 0.15 | 217 | U[217, 271]=42,715, p<0.0001**, r=0.39 |
| Q7. On a scale of 1-5 (1 most useful, 5 least useful), how useful was the reviewer feedback in informing your future scientific endeavors in the proposed research area? | 3.0 ± 0.12 | 342 | 4.0 ± 0.16 | 228 | U[228, 342]=50,536, p<0.0001**, r=0.25 |

Effect size r is Z-score/ $\sqrt{N}$ . \*\* indicates p<0.01

## Discussion

### Perceptions of Grant Review Feedback

The results of our analysis suggest that while the majority of grant applicants in our survey deemed reviewer feedback to be appropriate across several dimensions – including how fair, well-written and well-informed the feedback was – there were sizable proportions (40%-44%) who did not find it appropriate. Similar to the 2017 NIH survey, we found respondent perceptions were influenced significantly by funding success, where funded applicants found the feedback to be more appropriate. Admittedly, this variability by funding status highlights the subjectivity inherent in applicant perceptions. Nevertheless, respondent applicants noted a variety of specific types of perceived bias in the feedback, including conservatism, methodology bias, and gender bias. In addition, reliability concerns intersected with views on fairness; several respondents noted significantly different sets of issues identified in the feedback from panel to panel for resubmissions, revealing important inconsistencies in feedback that can be construed as inequitable to the applicant. In addition, multiple respondents mentioned the apparent lack of expertise of reviewers in areas of science related to the proposal, experience with animal models, and statistics. Applicants also indicated specific instances where the format, length and quality of the writing was insufficient; some commented that it appeared the reviewer had not fully read the proposal, as they penalized applicants for issues in the feedback that were specifically addressed in the proposal. While this lack of attention or preparation may reflect the state of overburden experienced by many peer reviewers (Gallo et al. 2020a), it is clear that the lack of appropriate reviewer feedback is still an issue for many applicants. However, this interpretation could be slanted, as respondents with comments generally had more negative views of the appropriateness of feedback as compared to non-commenting respondents.

Importantly, respondent perception of appropriate feedback differed by race and gender. Non-white respondentss found their feedback particularly unfair, suggest a potential perception of racial biases. This finding is in line with previous findings that suggest URM women are much less likely to resubmit an unfunded application compared to white women or men (Ginther et al. 2016); racially biased feedback could be contributing to these low resubmission rates, which in turn could contribute to current funding gaps among underrepresented groups (Ginther et al. 2011). These results could also suggest that there are significant perceived biases at play in the peer review process, which have also been suggested by scoring data (Tamblyn et al. 2018), and critique analysis (Pier et al. 2017). We also found that women were more likely to perceive their feedback as well written, cohesive, and balanced than men, but no gender difference was found for perceptions of usefulness of the feedback for preparing grant resubmissions. These results may differ from studies that found women to be more sensitive to peer feedback than men (Mayo 2016, Roberts & Nolen-Hoeksema, 1989), to have inaccurate negative self-perceptions about the quality of their work (Beyer 1998), and to have less motivation to resubmit unfunded applications (Biernat et al. 2020). A re-examination of racial and gender bias in the peer review process is needed as is the testing and application of valid mitigation strategies to reduce bias in review panels. Impartiality is needed to ensure the legitimacy of the peer review system, because the outcomes of such review panels are linked to funding and career trajectories and contribute to funding disparities currently seen among racial groups (Erosheva et al., 2020). Thus, ensuring fair reviews should help improve diversity in the scientific community, which has been shown to promote innovation (Hofstra et al. 2020).

Our analysis also indicates that the majority of our respondents did not find review feedback very useful in improving grantsmanship, future submissions, or future areas of research. These results may hint at a breakdown in one of the secondary functions of peer review––providing useful feedback to applicants; although funding agencies vary in their expectations for grant reviewers to provide such feedback. Interestingly, perceptions of usefulness were quite negative independent of applicant funding status, and the demographics of the respondents did not appear to significantly affect perceptions. Some respondents listed the format of the feedback and the lack of constructive criticism as issues, although some felt comments were relevant and potentially useful. Many comments focused on the low funding success rates and the low reliability between reviewers, and between original reviews and those of resubmissions as important hurdles to the utility of feedback (e.g., one can address initial reviewer concerns, only to have a separate set of concerns appear in the review of the resubmission). This reported lack of reliability may highlight some issues with the structure of the review process for resubmissions, which are exacerbated by poor funding rates. Overall, more effort should be placed on ensuring that the process of providing feedback is strengthened to achieve all goals, primary and secondary, of the peer review of applications.

Some respondents also mentioned how the appropriateness of feedback limits its usefulness for future submissions and our analysis confirms that negative perceptions of appropriateness are correlated with negative perceptions of usefulness. This relationship between appropriateness and usefulness is particularly concerning given the racial and gender differences found in the appropriateness ratings. While racial/gender factors do not seem to influence perceptions of the usefulness of feedback directly in our study, respondents may perceive different reasons for the lack of utility, and the perception of persistent bias in the funding system may be a strong influence on scientists’ decisions to submit projects for funding, or even to continue on a research career track. These effects should be examined more rigorously by funding agencies (and the results made public and prominent) to ensure an equitable review process and a healthy, diverse set of applicants.

### Generalizability

Our results are limited by a relatively low response rate, although this response rate approximates the rate of similar surveys on peer review (Gallo et al. 2020a, Ware 2008). The majority of funding agency comments in our survey mentioned the NIH as a recent source of review feedback; the gender, race, and degrees of our sample are similar to those reported from surveys and analyses of NIH applicants (NIH, 2012a, Ginther 2011, Ginther 2016). Our respondents tended to be older than NIH applicants (NIH, 2012a) but comparable in age to funded NIH investigators (Daniels 2015), consistent with our sample’s high funding success rate.

Despite the similarity of our overall sample to current applicant pools, white respondents were overrepresented among those who made comments. Future studies should include larger samples of underrepresented minorities to better examine differences between racial/ethnic groups and the motivations for applying or not applying. These results should be also replicated for groups of investigators whose applications were reviewed on the same panel, such that differences across scientific topic areas and funding mechanisms are minimized. Finally, an important limitation of this study is the potential effects of funding agency, as this variable was not assessed in our survey. While the majority of our respondents are likely referring to the NIH process based on their comments, some referred to NSF, which has a different review process. Future studies should include this factor in their analysis.

### Conclusion

In conclusion, our results suggest more emphasis should be placed on training reviewers to create constructive, useful, and appropriate feedback and enhancing procedures that detect strong biases and inappropriate comments early in the process, before they influence funding decisions and are communicated to applicants. These recommendations are in line with those from a 2012 report from the NIH Working Group on Diversity in the Biomedical Research Workforce, which was formed in response to reports of racial funding disparities (NIH 2012b). Our results also may suggest more progress on these recommendations is needed to “clarify the root causes for funding disparities” and to “significantly support the development and evaluation of programs that will increase diversity in the biomedical workforce”

## Appendix

**Supplementary Table 1.** Respondent Comments Grouped by Text Presence of Keyword Relevant to Survey Question

| Keyword category (Relevant Survey Question) | % of Total Grouped Comments (N) | % Positive Only | %Negative Only | %Both Pos/Neg |
| --- | --- | --- | --- | --- |
| Written<br>*Q1: Did you feel the reviewer feedback was well written, cohesive, and balanced?* | 10% (8) | 0% | 38% | 63% |
| Bias<br>*Q2: Did you feel the reviewer feedback was fair and unbiased?* | 33% (26) | 0% | 54% | 46% |
| Expertise<br>*Q3: Based on the reviewer feedback you received, do you feel that the reviewers had the appropriate expertise to evaluate your grant application?* | 48% (38) | 0% | 39% | 61% |
| Useful/Helpful<br>*Q4. On a scale of 1-5 (1 most useful, 5 least useful), overall how useful was the reviewer feedback you received on your last grant submission?*<br>*Q5. On a scale of 1-5 (1 most useful, 5 least useful), how useful was the reviewer feedback in improving your grantsmanship?*<br>*Q6. If you were not funded, on a scale of 1-5 (1 most useful, 5 least useful), how useful was the reviewer feedback in improving your future submissions?*<br>*Q7. On a scale of 1-5 (1 most useful, 5 least useful), how useful was the reviewer feedback in informing your future scientific endeavors in the proposed research area?* | 33% (26) | 8% | 31% | 62% |

Proportions calculated based on a total of 79 comments

**Supplementary Table 2.**
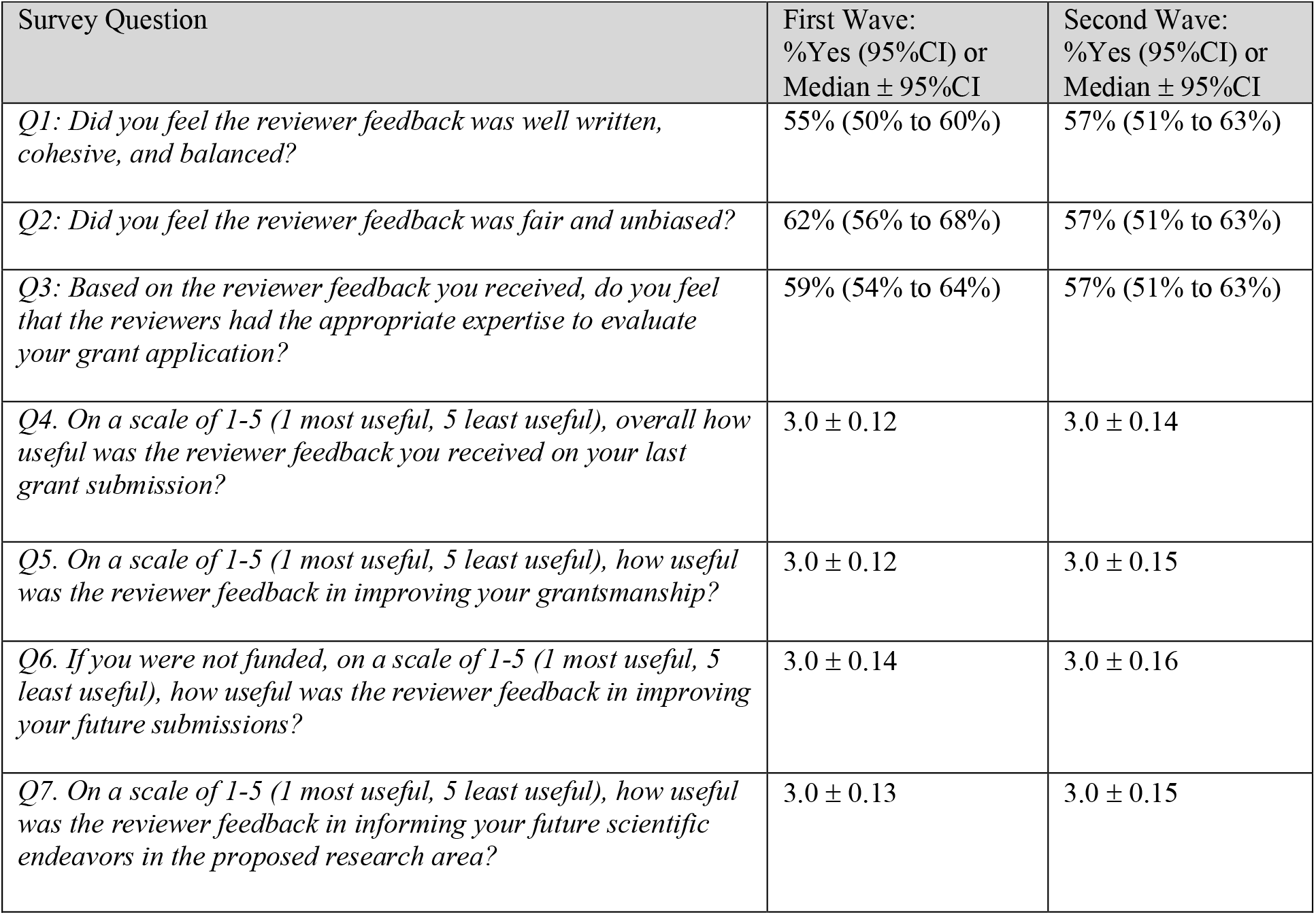
Comparisons of the quantitative answers to questions for Usefulness and Appropriateness for two waves of responses (first wave versus second wave).

**Supplementary Table 3.**
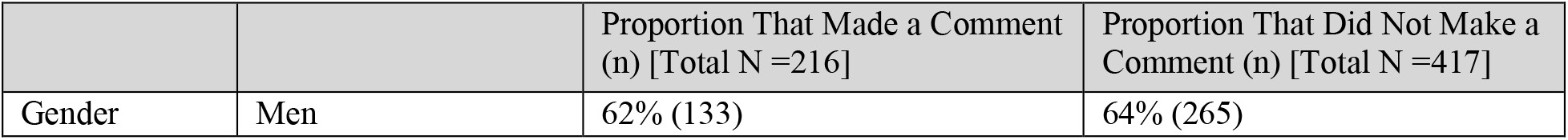
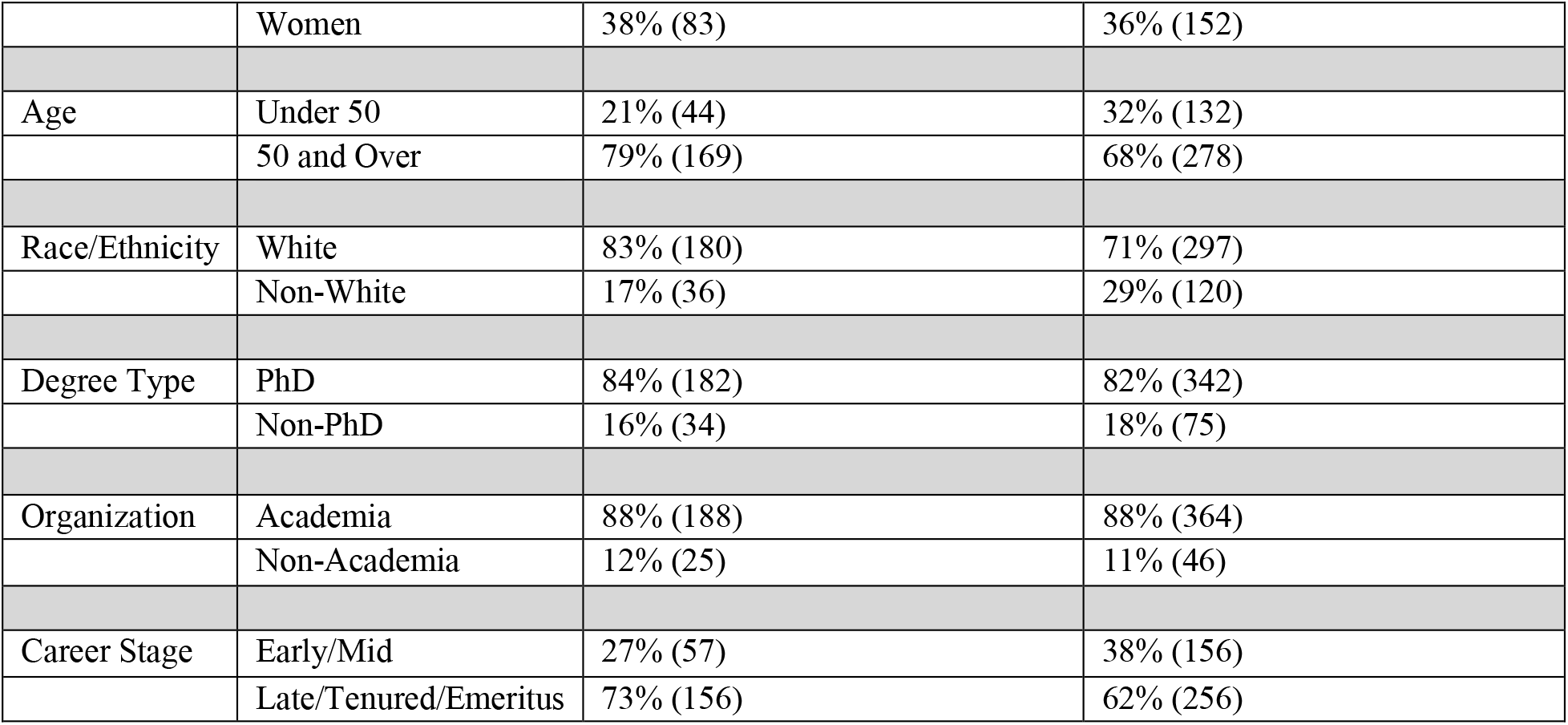
Demographics and Respondents with Comments

**Supplementary Table 4.** Example Respondent Comments Grouped by Survey Question

| Relevant Survey Question | Number | Comment |
| --- | --- | --- |
| *Q1: (well written, cohesive, and balanced?)* | Q1.1 | *“Current reviews are very short and written as bullet points. Therefore, there is normally criticism but no guidance which makes the review absolutely not useful for future improvement/applications.”* |
|  | Q1.2 | *“The critiques seemed to be hastily written (as they are, from my live-panel reviewer experience). Most annoyingly, NIH reviewers often down-score a proposal without comment, although again from my experience as a reviewer, I know they are instructed NOT to do that. In such cases they are in essence telling me that there was something they didn't like, but couldn't or wouldn't say what it was. This makes the information un-actionable, and thus useless.”* |
| *Q2: (fair and unbiased?)* | Q2.1 | *“Reviewers these days are often quite biased towards specific methodologies, often the ones they use.”* |
|  | Q2.2 | *“There is too much personal bias in grant review. Reviewers seem to have the people they want to champion and shoot down others they do not know.”* |
|  | Q2.3 | *“I believe there is too much in the way of politics and also bias against women in the peer review process.”* |
|  | Q2.4 | *“It takes just one biased or not knowledgeable reviewer to sink a grant application.”* |
| Q3: (appropriate expertise?) | Q3.1 | *“I feel that reviewers often lack technical background in my field and instead substitute comments related to empirical details that should be less important. But some reviewers do possess the necessary technical background.”* |
|  | Q3.2 | *“The last grant submitted was a mix of oncology, nutrition and immunology. No one reviewer could cover all of this expertise.”* |
| Q4-7: (useful reviewer feedback?) | Q4-7.1 | *“The last review was very short, but it did touch on some very important questions about the feasibility of completing the project. It is worth adding that this was a project that was previously submitted twice before to different organizations. Each time, the reviews were incredibly helpful and I believe they led me to improve the proposal during each iteration to the point it was eventually funded... which is the whole point of the peer review... constructive criticism.”* |
|  | Q4-7.2 | *“The application scored on the verge of fundability, this score had clearly been lowered by people who had voted outside the reviewer’s limits. The resubmission, incorporating improvements suggested by the original review, was triaged. Such inconsistency between panels implies a profoundly unreliable quality of review and tends to confirm the view that we are in a casino rather than an effectively regulated system.”* |
|  | Q4-7.3 | *“While reviewers have often been helpful, they are frequently less helpful when they make sweeping comments outside their area of expertise, especially statistics – where they sometimes criticize methods they are unfamiliar with or don’t understand, or worse- yet, ask for irrelevant/inappropriate analyses to be added.”* |
|  | Q4-7.4 | *“The reviews were one sentence ‘bullets’ (these were NIH applications). NIH has (in my experience as a reviewer within the past decade) given reviewers explicit instructions to NOT offer suggestions for alternative approaches – and the reviews we received stated only that ‘X was not likely to be useful / informative / etc.’.”* |
|  | Q4-7.5 | *“The reviews are often helpful, but frequently they are also misfires. By that, I mean they bring up some issue that nobody else has ever brought up, and something we will never see again. If we transform our proposal in response to such unusual comments, we will be bouncing around too much. Yet such a comment can kill a grant. There is far too much noise in the system. Also, the ability to distinguish good grants from excellent ones is diminished because of such noise. Say that adds +/-10% error into the system. Yet the pay lines are sometimes at this level of noise!”* |

**Supplementary Table 5.**
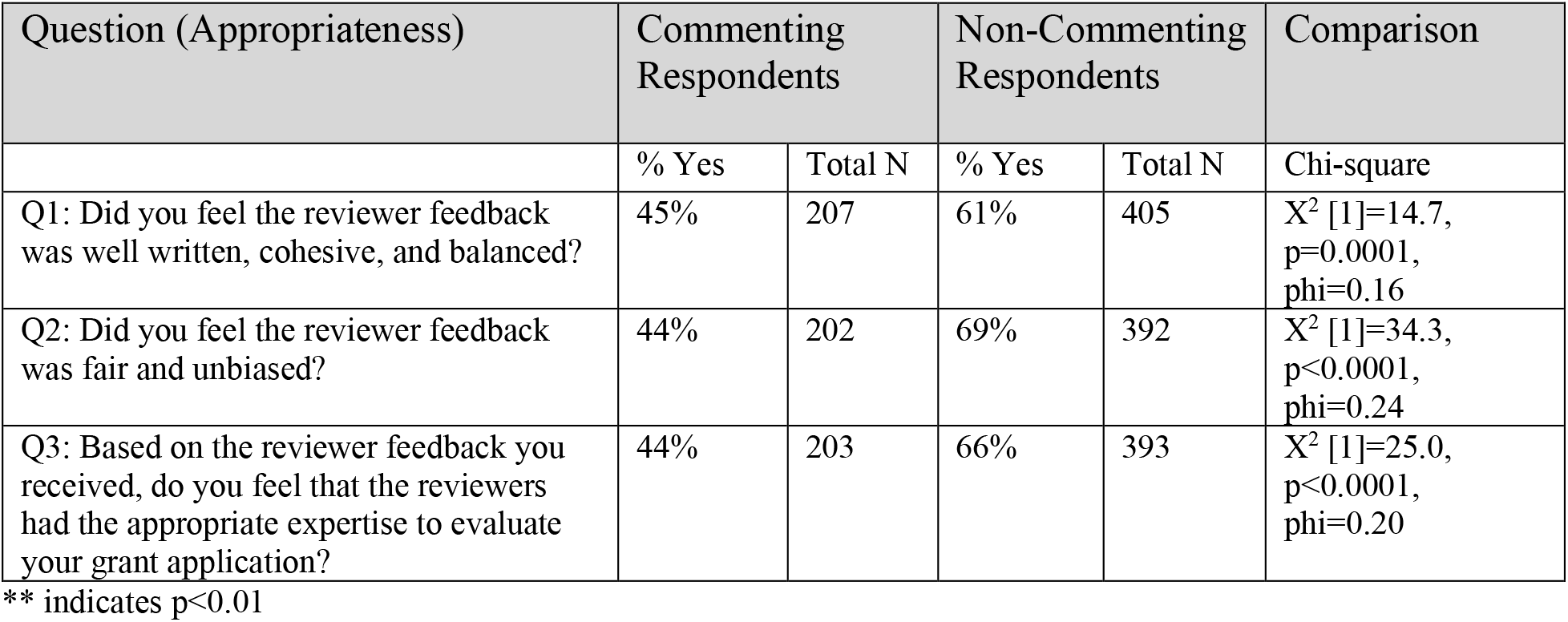
Answers for Appropriateness Questions and Respondents with Comments

**Supplementary Table 6.** Variance Inflation Factors (VIFs) for independent variables from the logistic regression models for Q1-7

| Factor | Men | Age (Over 50)? | White? | PhD? | Academia? | Early / Mid-Career? | Funded? |
| --- | --- | --- | --- | --- | --- | --- | --- |
| VIF – Q1 (Well Written) | 1.02 | 1.29 | 1.04 | 1.02 | 1.02 | 1.33 | 1.01 |
| VIF – Q2 (Fair/Unbiased) | 1.02 | 1.29 | 1.03 | 1.02 | 1.02 | 1.31 | 1.01 |
| VIF – Q3 (Expertise) | 1.02 | 1.34 | 1.04 | 1.03 | 1.03 | 1.37 | 1.01 |
| VIF – Q4 (Overall Useful) | 1.03 | 1.33 | 1.03 | 1.03 | 1.03 | 1.36 | 1.01 |
| VIF – Q4 (Grantsmanship-Useful) | 1.03 | 1.35 | 1.05 | 1.03 | 1.04 | 1.39 | 1.01 |
| VIF – Q4 (Future Submissions-Useful) | 1.04 | 1.32 | 1.05 | 1.03 | 1.03 | 1.33 | 1.02 |
| VIF – Q4 (Future Research Endeavors-Useful) | 1.03 | 1.30 | 1.05 | 1.02 | 1.04 | 1.34 | 1.01 |

## Declarations

### Ethics approval

The study was reviewed by the Washington State University Office of Research Assurances (Assurance# FWA00002946) and granted an exemption from IRB review consistent with 45 CFR 46.101(b)(2).

### Consent to participate

Participants were free to choose whether or not to participate in the survey and consented by their participation.

### Consent for publication

Not applicable.

### Availability of data and material

A full copy of the peer review survey is available in the **S1 File** in the Supporting Information. The raw, anonymized data are available on Figshare (https://doi.org/10.6084/m9.figshare.8132453.v1).

### Authors’ contributions

SAG, KBS, LAT, and SRG had full access to all the data in the study and take responsibility for the integrity of the data and the accuracy of the data analysis. SAG, KBS, and LAT contributed to study concept and design; KBS prepared the Institutional Review Board application and was its point of contact; LAT implemented the survey data collection; SAG and LAT exported and prepared the data; all authors contributed to the analysis and interpretation of the data; SAG drafted the manuscript, KBS, LAT and SRG contributed to the critical revision of the manuscript; and SAG provided study supervision. All authors read and approved the final manuscript.

## Supporting Information

**S1 File. Copy of Full Survey Text**

Survey is listed below.

[]What is your gender?

Please choose all that apply:

Male

Female

Prefer not to answer

[]What is your age?

Please choose only one of the following:

Under 30

30–39

40–49

50–59

60+

[]Please specify your race/ethnicity

Please choose all that apply:

American Indian or Alaska Native

Asian or Asian American

Black or African American

Hawaiian or Other Pacific Islander

Hispanic or Latino

Non-Hispanic White/Caucasian

Other

Prefer not to answer

[]What type of degree(s) do you have?

Please choose all that apply:

PhD or other research doctorate

MD

DDS

DVM or VMD

Other

Prefer not to answer

[]What type of an organization do you work for?

Please choose only one of the following:

Academia

Government

Industry

Other

[]What stage of career have you reached?

Please choose only one of the following:

Early career

Mid career

Late career/tenured

Emeritus

[]On average, how many hours do you work each week?

Please choose only one of the following:

40 h

40–50 h

50–60 h

60–70 h

70 + h

[]Please provide any comments that justify your responses under Section 1, Demographics. Please write your answer here:

Section 2: Grant submission and peer review experience

[]Have you submitted a grant for peer review in the last 3 years?

Please choose only one of the following:

Yes

No

[]If you answered yes to submitting a grant for peer review in the past 3 years, how many grant applications have you submitted in that time frame?

Please choose only one of the following:

1

2

3

4

5

6

7 or more

[]Did you receive reviewer feedback on your last grant submission?

Please choose only one of the following:

Yes

No

[]Was your last application successful, i.e., were you funded?

Please choose only one of the following:

Yes

No

[]Have you served on a peer review panel in the last 3 years?

Please choose only one of the following:

Yes

No

[]If you answered yes to serving on a peer review panel in the last 3 years, how many peer review panels have you served on in that time frame?

Please choose only one of the following:

1

2

3

4

5

6

7 or more

[]If you answered yes to serving on a peer review panel in the past 3 years, please select the mode of your last peer review panel meeting.

Please choose only one of the following:

Face-to-face

Remote (video/teleconference)

Internet-assisted

Other

[]How many ad-hoc reviews (usually one or two grant applications reviewed telephonically that are being evaluated in a panel meeting setting) have you performed in the past 3 years?

Please choose only one of the following:

0

1

2

3

4

5

6

7 or more

[]Have you reviewed for a journal in the last 3 years?

Please choose only one of the following:

Yes

No

[]If you answered yes to reviewing for a journal in the past 3 years, how many submissions have you reviewed in that time frame?

Please choose only one of the following:

1

2

3

4

5

6

7 or more

[]What is a higher personal priority: grant review or journal review?

Please choose only one of the following:

Grant review

Journal review

Both are equal priority

Neither is a priority

[]Please elaborate on your responses under Section 2, Grant submission and peer review experience.

Please write your answer here:

Section 3: Investigator attitudes toward grant review

If you answered yes to receiving feedback on your last grant submission, please answer Section 3 of the questionnaire. If you answered no, please proceed to Section 4.

[]On a scale of 1–5 (1 most useful, 5 least useful), overall how useful was the reviewer feedback you received on your last grant submission?

Please choose only one of the following:

1

2

3

4

5

[]On a scale of 1–5 (1 most useful, 5 least useful), how useful was the reviewer feedback in improving your grantsmanship?

Please choose only one of the following:

1

2

3

4

5

[]If you were not funded, on a scale of 1–5 (1 most useful, 5 least useful), how useful was the reviewer feedback in improving your future submissions?

Please choose only one of the following:

1

2

3

4

5

[]On a scale of 1–5 (1 most useful, 5 least useful), how useful was the reviewer feedback in informing your future scientific endeavors in the proposed research area?

Please choose only one of the following:

1

2

3

4

5

[]Did you feel the reviewer feedback was well written, cohesive, and balanced?

Please choose only one of the following:

Yes

No

[]Did you feel the reviewer feedback was fair and unbiased?

Please choose only one of the following:

Yes

No

[]Overall, in what area(s) did the reviewer feedback primarily focus?

Please choose all that apply:

Potential impact of research

Hypothesis

Research methodology

Innovation potential

Preliminary data

Responsiveness to funding mechanism

Statistical issues

Qualifications of research team

Budget

Other

[]Did the reviewers comment on the riskiness of the research project?

Please choose only one of the following:

Yes

No

[]Based on the reviewer feedback you received, do you feel that the reviewers had the appropriate expertise to evaluate your grant application?

Please choose only one of the following:

Yes

No

[]Please elaborate on your responses under Section 3, Investigator Attitudes Towards Grant review.

Please write your answer here:

Section 4: Reviewer attitudes towards grant review

[]What are your reasons for accepting an invitation to serve on a peer review panel?

Please choose all that apply:

Desire to give back to the scientific community

Networking opportunities

Informing your own grantsmanship

Gaining exposure to new and innovative scientific areas

Enhancing your career/resume

Expectation from the funding agency

Honorarium

Other

[]Do you feel that serving as a reviewer on peer review panels has positively impacted your career?

Please choose only one of the following:

Yes

No

[]If you feel that serving as a peer reviewer has positively impacted your career, in what ways has serving as a reviewer influenced your career?

Please choose all that apply:

Bolstered your career

Improved your grantsmanship

Increased your exposure to new scientific ideas

Improved your networking/collaboration opportunities

Other

[]In general, which type of panel meeting format do you prefer?

Please choose only one of the following:

Face-to-face

Virtual [teleconference/videoconference]

Internet-assisted

[]On a scale of 1–5, (1 most influential, 5 least influential), please rate the following factors in influencing your selection of preferred panel meeting format:

Please write your answer(s) here:

Logistical convenience

Level of communication among panel members

Networking opportunities

Likelihood to participate on panel

[]In the last 3 years, how many times have you declined an invitation to serve on a peer review panel?

Please choose only one of the following:

1

2

3

4

5

6

7 or more

[]What were your reasons for declining an invitation to serve on a peer review panel?

Please choose all that apply:

Limited free time

Poor expertise match

Personal reasons (holiday, sickness, travel)

Review timeline too compressed

Conflict of interest

Issue with funding agency

Other

[]What is the maximum number of peer review panels/committees you prefer to serve on per year?

Please choose only one of the following:

1

2

3

More than 3

[]What is the maximum number of days you prefer to attend a peer review panel meeting? Please choose only one of the following:

1

2

3

More than 3

[]What is the maximum number of R01-type grant applications you prefer to be assigned for a peer review panel meeting?

Please choose only one of the following: 1–2

3–4

5–6

7

More than 7

[]What was the actual number of days of your last peer review panel meeting?

Please choose only one of the following:

1

2

3

More than 3

[]What was the actual number of R01-type grant applications you were assigned to review at your last peer review panel meeting?

Please choose only one of the following:

1–2

3–4

5–6

7–8

More than 8

[]On average, how many hours did you spend reviewing each grant application before the panel meeting?

Please choose only one of the following:

1–2

2–3

3–4

4–5

5–6

7 or more

[]Please elaborate on your responses under Sect. 4, Reviewer attitudes towards grant review. Please write your answer here:

Section 5: Peer review panel meeting proceedings

[]Please answer the following questions in relation to your last peer review meeting. On a scale of 1–5 (1 most definitely, 5 not at all), was your scientific expertise necessary and appropriately used in the review process?

Please choose only one of the following:

1

2

3

4

5

[]On a scale of 1–5 (1 most definitely, 5 not at all), from your perspective was the expertise of the other panel members necessary and appropriately used in the review process?

Please choose only one of the following:

1

2

3

4

5

[]Did the grant application discussions facilitate reviewer participation?

Please choose only one of the following:

Yes

No

[]Were the grant application discussions fair and balanced?

Please choose only one of the following:

Yes

No

[]On a scale of 1–5 (1 most useful, 5 least useful), how useful were the grant application discussions in clarifying differing reviewer opinions?

Please choose only one of the following:

1

2

3

4

5

[]On a scale of 1–5 (1 extremely effective, 5 no effect), did the grant application discussions affect the outcome?

Please choose only one of the following:

1

2

3

4

5

[]On a scale of 1–5 (1 most appropriate, 5 least appropriate), were the evaluation criteria appropriate to judge the best science and move the field forward?

Please choose only one of the following:

1

2

3

4

5

[]On a scale of 1–5 (1 extremely important, 5 of no importance), how important is the PI’s track record to assessing an investigator initiated (R01)-type application?

Please choose only one of the following:

1

2

3

4

5

[]In general, does a PI’s track record temper your assessment of any detected methodological weaknesses?

Please choose only one of the following:

Yes

No

[]On a scale of 1–5 (1 most definitely, 5 not at all), did the grant application discussions promote the best science?

Please choose only one of the following:

1

2

3

4

5

[]Was innovation factored into selecting the best science?

Please choose only one of the following:

Yes

No

[]Did you view innovation as an essential component of scientific excellence when evaluating the grant applications?

Please choose only one of the following:

Yes

No

[]Did the risks associated with innovative research impact the scores you assigned to the grant applications?

Please choose only one of the following:

Yes

No

[]On a scale of 1–5 (1 completely, 5 not at all), how much did the seniority of your fellow panel members influence your evaluations during the panel deliberations?

Please choose only one of the following:

1

2

3

4

5

[]Was the format and duration of the grant application discussions sufficient to allow the non-assigned reviewers to cast well informed merit scores?

Please choose only one of the following:

Yes No

[]On a scale of 1–5 (1 extremely useful, 5 not useful at all), how useful was the Chair in facilitating the application discussions?

Please choose only one of the following:

1

2

3

4

5

[]Please elaborate on your responses under Section 5, Peer review panel meeting proceedings. Please write your answer here:

Thank you for taking the time to fill out the survey. Have a wonderful day!

Submit your survey.

Thank you for completing this survey.

